# Removal of the Ciliary Gate Allows Axoneme Extension in the Absence of Retrograde IFT

**DOI:** 10.1101/2025.10.21.683645

**Authors:** Ana R. G. De-Castro, Maria J. G. De-Castro, Guus H. Haasnoot, Reto Gassmann, Erwin J. G. Peterman, Tiago J. Dantas, Carla M. C. Abreu

**Affiliations:** i3S - Instituto de Investigação e Inovação em Saúde, Universidade do Porto, Porto, Portugal; IBMC - Instituto de Biologia Molecular e Celular, Universidade do Porto, Porto, Portugal; ICBAS - Instituto de Ciências Biomédicas Abel Salazar, Universidade do Porto, Porto, Portugal; Department of Physics and Astronomy and LaserLaB, Vrije Universiteit Amsterdam, Amsterdam, Netherlands; Life and Health Sciences Research Institute (ICVS), School of Medicine, University of Minho, Braga, Portugal; ICVS/3B’s-PT Government Associate Laboratory, Braga, Portugal

**Keywords:** cilium, intraflagellar transport, retrograde transport, ciliary extracellular vesicles, diffusion barrier

## Abstract

The transition zone (TZ) is a selective barrier that maintains ciliary compartmentalization by controlling protein entry and exit. Cilia assembly requires the crossing of this barrier by intraflagellar transport (IFT) trains, scaffolded by IFT-A and IFT-B complexes, which move cargo bidirectionally using kinesin-2 and dynein-2 motors.

In *Caenorhabditis elegans*, IFT-A loss abolishes retrograde transport, resulting in truncated cilia packed with IFT material. Here, we show that blocking TZ assembly prevents dynein-2 and IFT-B accumulation inside IFT-A-deficient cilia and partially rescues axoneme length. Single-particle imaging reveals that this rescue occurs without recovery of retrograde IFT. Instead, IFT particles exit cilia by passively diffusing through the disrupted TZ. Moreover, IFT-A/TZ double mutants shed ciliary extracellular vesicles (EVs) abnormally enriched in IFT components, providing a second clearance route. We conclude that TZ removal alters ciliary responses to retrograde transport defects, promoting diffusion and EV release to clear IFT machinery and facilitate axoneme extension.

**Highlights:**

- TZ loss provides alternative routes for clearing IFT machinery stalled in IFT-A mutant cilia
- Axoneme extension is possible without retrograde IFT when the TZ barrier is removed
- Disrupted TZ enables exit of IFT particles by passive diffusion in retrograde IFT-deficient cilia
- Excess IFT machinery is discarded in ciliary EVs when retrograde IFT and gating are compromised

## INTRODUCTION

Cilia are microtubule-based organelles that project from the surface of many eukaryotic cells, playing crucial roles in cell signaling, fluid movement, and sensory perception. Defects in their assembly or function cause ciliopathies, severe congenital disorders that impact multiple organ systems (Mill et al., 2023; Reiter and Leroux, 2017). The assembly and maintenance of cilia rely on a highly conserved process known as intraflagellar transport (IFT)(Kozminski et al., 1993), which moves molecular cargo along the microtubules of the ciliary axoneme.

This bidirectional transport of ciliary cargos is mediated by large polymeric platforms, known as IFT trains, and the coordinated action of two opposing molecular motors: kinesins-2 drive anterograde transport from the base to the tip, while dynein-2 powers retrograde transport from the tip back to the base.

The primary scaffolds of IFT trains are the IFT-B and IFT-A protein complexes, which not only define train architecture but also provide binding sites for motors and adaptor complexes that mediate cargo binding (Lacey and Pigino, 2025). Notably, the IFT-A complex has been particularly implicated in the import of transmembrane and membrane-associated proteins into cilia (Reddy Palicharla and Mukhopadhyay, 2024).

The IFT-A complex can be subdivided into a core A1 module (IFT144, IFT140 and the C-terminal (CT) region of IFT122) and a peripheral A2 module (IFT121, IFT139, IFT43 and the N-terminus (NT) of IFT122) (Hesketh et al., 2022). IFT-A1 subunits are essential for IFT-A complex assembly and its incorporation into cilia (Goncalves-Santos et al., 2023). Loss of IFT-A1 subunits severely disrupts retrograde IFT, resulting in short, swollen cilia packed with IFT and motor components, a phenotype closely resembling dynein-2 null mutants (Pazour et al., 1999; Piperno et al., 1998; Reddy Palicharla and Mukhopadhyay, 2024). Our recent work has shown that IFT-A promotes dynein-2 activation at the ciliary tip, couples this motor to IFT-B in retrograde trains, and enhances retrograde transport velocity (Goncalves-Santos et al., 2023).

In addition to IFT, a selective gate between the ciliary base and the axoneme, known as the transition zone (TZ), is key for maintaining ciliary compartmentalization/separation from the cytoplasm (Garcia-Gonzalo and Reiter, 2017; Moran et al., 2024). This barrier is composed of large protein assemblies that are typically subdivided into the MKS and NPHP modules. Non-core IFT-A subunits and the dynein-2 motor itself have been implicated in supporting TZ assembly and barrier integrity (Jensen et al., 2018; Scheidel and Blacque, 2018). Notably, disruption of the IFT-140 core subunit, which abolishes the entire IFT-A complex, has only a relatively minor impact in TZ maintenance (Picariello et al., 2019; Scheidel and Blacque, 2018).

In *C. elegans*, the recruitment and assembly of both MKS and NPHP modules rely on MKS-5 (known as RPGRIP1L in vertebrates), which is the central organizer and scaffold of the TZ gate (Jensen et al., 2015; Williams et al., 2011). We have previously shown that disrupting the entire TZ via *mks-5* deletion or just the NPHP module via *nphp-4* deletion clears IFT particle accumulations caused by a reduction in dynein-2 motors, implicating the TZ as a physical gate that impedes the exit of IFT material when retrograde transport is weakened (De-Castro et al., 2022).

Emerging evidence shows that cilia composition can also be controlled by the shedding of small membrane-bound vesicles, known as ciliary extracellular vesicles (EVs)(Wang et al., 2014; Wood et al., 2013). These ciliary EVs typically carry signaling molecules and membrane proteins, and are implicated in inter-cellular/-animal communication and turnover of membrane components (Akella et al., 2020; Nager et al., 2017; Razzauti and Laurent, 2021; Salinas et al., 2017; Wang et al., 2024). Notably, under certain stress conditions, EV cargo composition can change, potentially enabling the export of other ciliary proteins from the cilium, including accumulated IFT subunits (Lobo et al., 2025; Nakamura et al., 2020; Phua et al., 2017)).

Here, we take advantage of the genetically tractable *C. elegans* model with well-defined ciliary architecture (Nechipurenko and Sengupta, 2025) to study how the interplay between IFT and ciliary gating impacts cilia homeostasis in neurons of a live animal. More specifically, we tested whether permeabilizing or eliminating the TZ barrier could alleviate the severe IFT particle congestion in IFT-A-deficient cilia. Remarkably, we find that *mks-5* deletion in *ift-140* or *ift-122* mutants (also known as *che-11* and *daf-10*) markedly reduces ciliary accumulations of dynein-2 and IFT-B components, and partially restores axoneme length. Live imaging reveals that this rescue occurs without recovery of active retrograde transport: instead, IFT particles exit by diffusion through the disrupted barrier. In addition, the combined loss of IFT-A and the TZ triggers extensive ectosome release carrying IFT machinery, suggesting that it acts as a second clearance route that complements diffusion. These results show that when retrograde IFT is abolished, removal of the TZ barrier enables passive diffusion and ectosome shedding as alternative pathways that clear IFT material and facilitate axoneme elongation.

## RESULTS

### Removing the TZ scaffold MKS-5 clears IFT accumulations and facilitates axoneme extension in IFT-A mutant cilia

In a prior study, we showed that loss of the dynein-2 intermediate chain, WDR-60, leads to the accumulation of IFT components inside cilia, at the distal side of the TZ (De-Castro et al., 2022). Disrupting the TZ in this mutant, either by deleting NPHP-4 or MKS-5, was sufficient to relieve these accumulations, indicating that the TZ offers resistance to the exit of retrograde IFT trains when dynein-2 is weakened. Therefore, we investigated whether disrupting TZ components would impact the very severe accumulations of IFT particles and dynein-2 characteristic of IFT-A1 mutants. We focused on *ift-140(tm3433)* and *ift-122(tm2878)*, two deletion alleles that completely destabilize the IFT-A complex, and produce short, swollen cilia, overloaded with IFT components as in dynein-2 *null* mutants (Goncalves-Santos et al., 2023).

Unlike in NPHP-4/WDR-60 mutants (De-Castro et al., 2022), disrupting the NPHP module using the *nphp-4(tm925)* allele in IFT-A mutant backgrounds did not reduce dynein-2 accumulations; in fact, approximately one-third of cilia contained even higher levels of dynein-2 inside, suggesting aggravated retention (Figure S1). In contrast, combining either IFT-A mutant with the *mks-5(tm3100)* allele, which completely disrupts the TZ barrier (Jensen et al., 2015; Williams et al., 2011), resulted in a marked reduction in the levels of dynein-2 motor subunits DHC-2 (also known as CHE-3) and WDR-60 inside cilia (Figures 1A-C).

**Figure 1.**
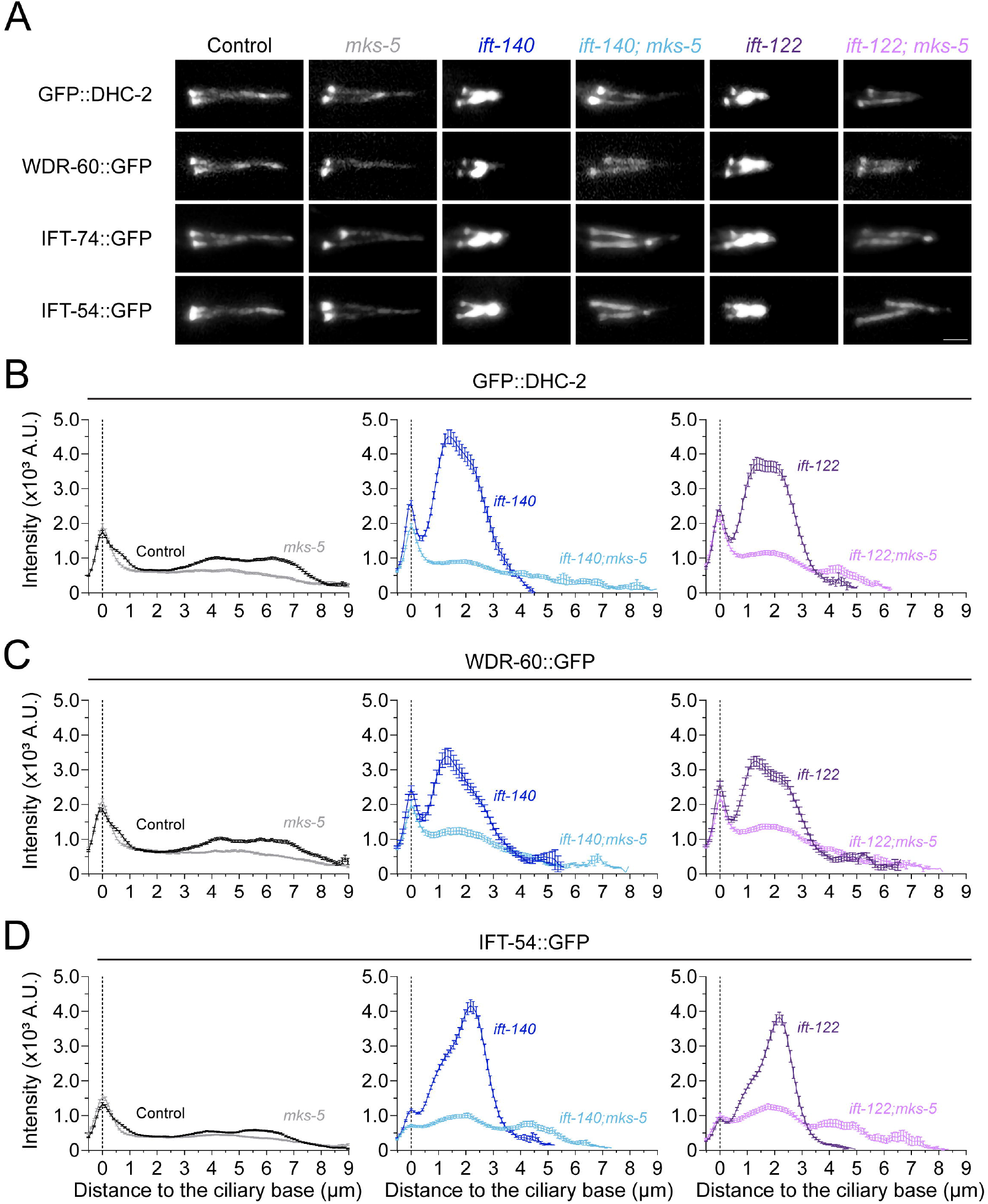
TZ removal clears IFT particle accumulations in IFT-A-deficient cilia. **(A)** Phasmid cilia from control, IFT-A, *mks-5* single and double mutant animals expressing endogenously tagged IFT components: dynein-2 heavy chain (GFP::DHC-2, also known as GFP::CHE-3), dynein-2 intermediate chain (WDR-60::GFP), IFT-B1 subunit (IFT-74::GFP), and IFT-B2 subunit (IFT-54::GFP, also known as DYF-11::GFP). Scale bar, 2 μm. **(B-D)** Distribution profile of GFP-tagged DHC-2 (B), WDR-60 (C), and IFT-54 (D) along cilia of the indicated genotypes (n ≥ 50 cilia per condition tested). The base of cilia is set at the position 0 (dotted line). Data represent mean ± SEM.

Given our previous finding that IFT-A is essential for coupling IFT-B to dynein-2 during retrograde transport (Goncalves-Santos et al., 2023), we anticipated that IFT-B components would remain trapped in *ift-122;mks-5* or *ift-140;mks-5* double mutants (hereafter also referred to as IFT-A/TZ double mutants). Surprisingly, however, similar to dynein-2, IFT-54 (also known as DYF-11) and IFT-74 subunits of the IFT-B complex were also cleared from IFT-A-deficient cilia when the TZ was removed (Figures 1A and 1D).

Another striking observation was that TZ loss led to a significant increase in the ciliary length in both IFT-A mutant backgrounds (Figures 1A and 2B). Using TBB-4::eGFP, a well-established axoneme marker (Hao et al., 2011), and SPD-5::RFP to mark the ciliary base (Magescas et al., 2021), we confirmed that this indeed reflected a true increase in axoneme elongation in IFT-A mutant cilia upon MSK-5 deletion (Figure 2A and 2C).

**Figure 2.**
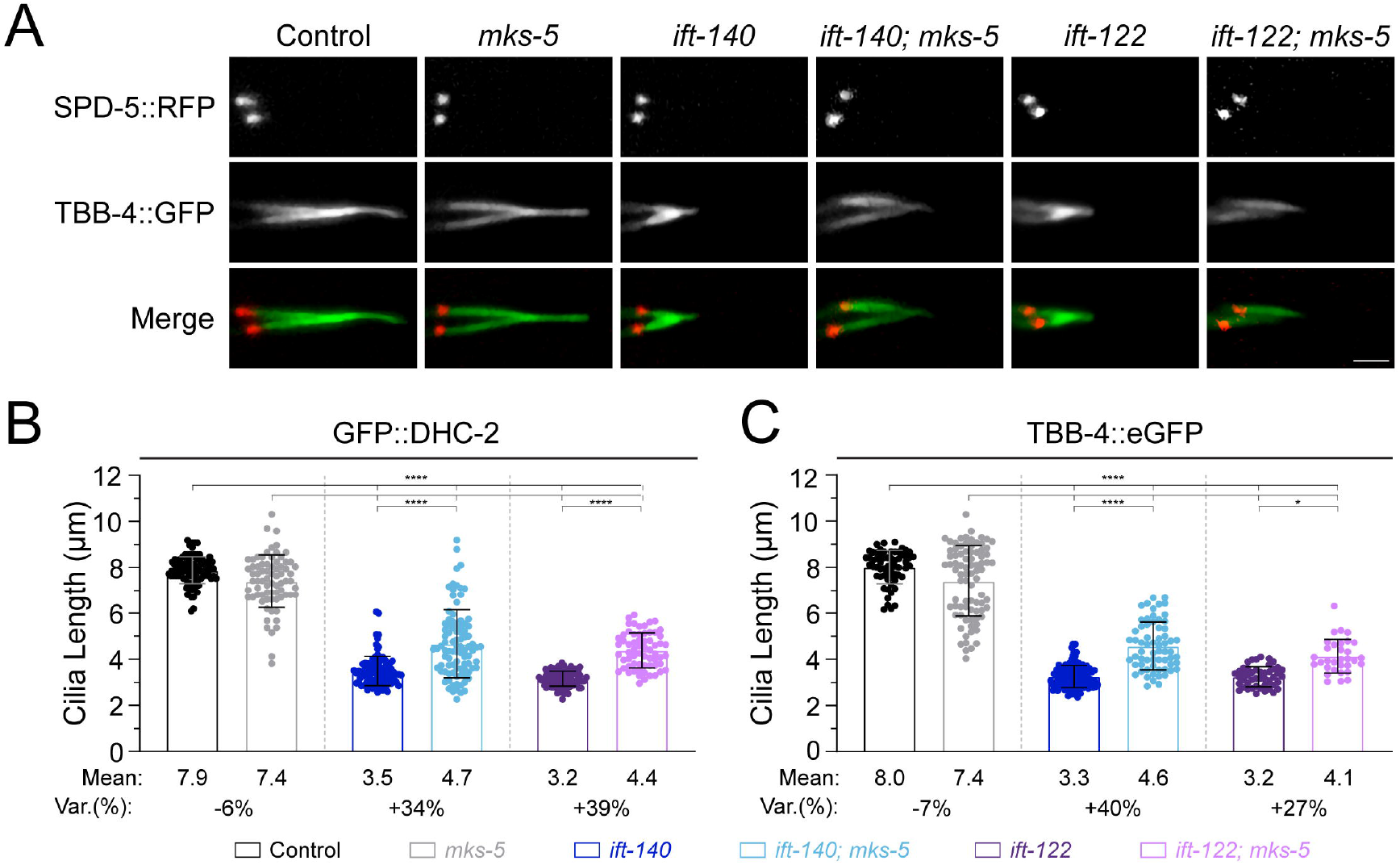
Disrupting the TZ facilitates axoneme extension in core IFT-A mutants. **(A)** Representative images showing axonemal microtubules (TBB-4::eGFP) and the ciliary basal body (SPD-5::RFP) in phasmid cilia from control, IFT-A, *mks-5* single and double mutant animals. Scale bars, 2 μm. **(B, C)** Quantification of ciliary length in control, IFT-A, *mks-5* single and double mutants expressing GFP::DHC-2 (B) or TBB-4::eGFP (C). One-way ANOVA followed by the Kruskal Wallis multiple comparisons test were used to analyze the datasets in B and C, respectively. *p ≤ 0.1; ****p ≤ 0.0001.

Together, these results suggest that loss of the TZ barrier allows the exit of IFT particles even when IFT train integrity is strongly impaired. Importantly, these findings also show that loss of the TZ barrier allows axoneme elongation in the absence of functional retrograde IFT.

### Disrupting the TZ gate enables diffusion-driven exit of IFT particles in IFT-A-deficient cilia despite impaired retrograde transport

To investigate how IFT particles exit the cilium in the combined absence of IFT-A and TZ, we performed live imaging of GFP::DHC-2 to observe IFT dynamics in single and double mutants.

Consistent with previous studies, retrograde IFT was virtually absent in both IFT-A mutants, whereas anterograde tracks persisted. Moreover, removing IFT-A did not reduce average anterograde IFT velocity in either the wild-type or the *mks-5* mutant backgrounds (Figures 3A and 3B). In fact, anterograde IFT-A-deficient trains reached higher velocities earlier than in controls in the first 3µm (Figure S2).

**Figure 3.**
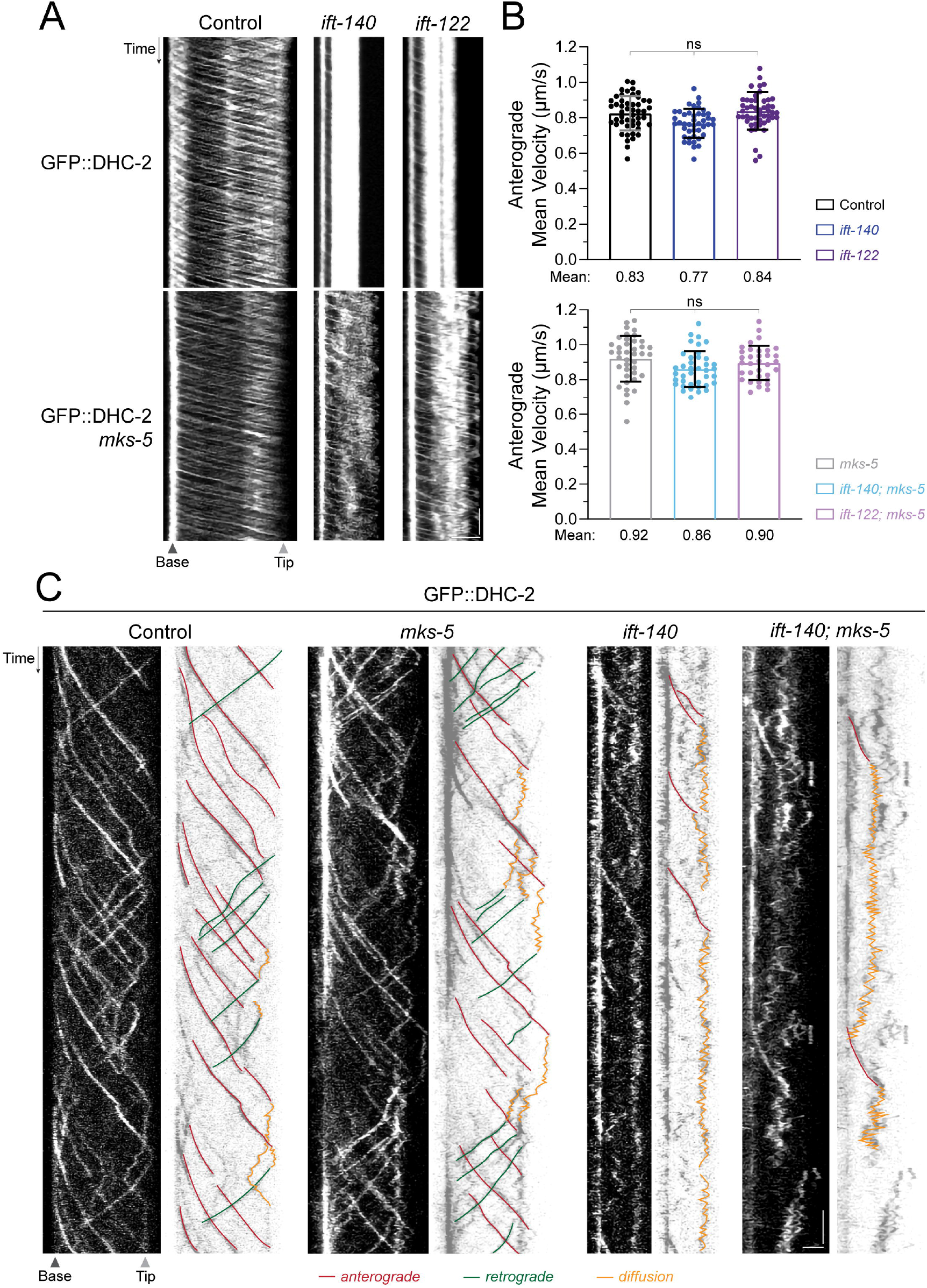
TZ loss preserves anterograde IFT and enables diffusion-based clearance of IFT particles in IFT-A-deficient cilia. **(A)** Kymographs of GFP::DHC-2 in control, IFT-A, *mks-5*, single and double mutant backgrounds obtained from live imaging. Scale bars: vertical, 5 s; horizontal, 2 μm. **(B)** Mean IFT velocities of GFP::DHC-2 control, IFT-A, *mks-5*, single and double mutants. Bars represent mean ± SD. One-way ANOVA followed by the Kruskal Wallis multiple comparisons test were used to analyze the datasets. ns, not significant. **(C)** Kymographs of GFP::DHC-2 in control, IFT-A, *mks-5*, single and double mutant backgrounds obtained using single particle fluorescence live imaging. Anterograde and retrograde particle trajectories are shown in red and green, respectively. Scale bars: vertical, 2 s; horizontal, 2 μm.

Disrupting the TZ via the *mks-5* mutation caused a modest but consistent increase in average anterograde IFT velocity across all backgrounds (Figure 3A and 3B, top panel vs. bottom panel), with trains reaching peak speeds earlier than in controls (Figure S2). This effect is consistent with previous reports that the TZ functions as a physical barrier, offering resistance to both the entry and exit of IFT trains and other ciliary proteins (De-Castro et al., 2022; Jensen et al., 2015; Prevo et al., 2015).

Retrograde IFT in IFT-A/TZ double mutants, on the other hand, was severely disrupted. Upon reaching the ciliary tip, their IFT trains failed to form the continuous, directed tracks characteristic of wild-type cilia. Instead, after turnaround, IFT particles dispersed slowly, moving in irregular, non-linear trajectories and forming a diffuse fluorescent “cloud”, a phenotype identical to that seen in IFT-A single mutants (Goncalves-Santos et al., 2023). Similar results were obtained when performing sequential photobleaching (Goncalves-Santos et al., 2023) to track isolated fractions of IFT particles (GFP::DHC-2) as they traveled along the ciliary axoneme of IFT-A and IFT-A/TZ double mutants (Figure S3A).

To resolve these heterogeneous particle behaviors, we employed high-resolution imaging with single-molecule sensitivity (Mitra et al., 2024; Zhang et al., 2021), which confirmed that returning GFP::DHC-2 particles in both IFT-A single and IFT-A/TZ double mutants exhibited non-processive, zig-zag motion rather than typical motor-directed, linear retrograde tracks (Figure 3C). Such irregular, non-linear movement has been associated with diffusion-like particle behavior in cilia (Mitra et al., 2024; Zhang et al., 2021). Importantly, while most diffusing particles in IFT-A single mutants remained trapped inside cilia, a substantial fraction in double mutants eventually exited through the disrupted TZ. A similar behavior was observed for IFT-B particles when carrying out high-resolution single-molecule imaging of IFT-54::GFP (Figure S3B).

Together, these findings indicate that removing the TZ does not restore active retrograde transport in IFT-A mutants but instead allows particle clearance through a diffusion-based mechanism.

### Combined loss of IFT-A and MKS-5 boosts release of ectosomes abnormally loaded with IFT machinery to relieve ciliary protein overload

During live imaging, we observed an additional, unexpected phenomenon in IFT-A/TZ double mutants: the presence GFP::DHC-2 and IFT-54::GFP in vesicle-like structures, which were frequently released from multiple sites along the ciliary membrane, including at the distal tip and base (Figure 4A).

**Figure 4.**
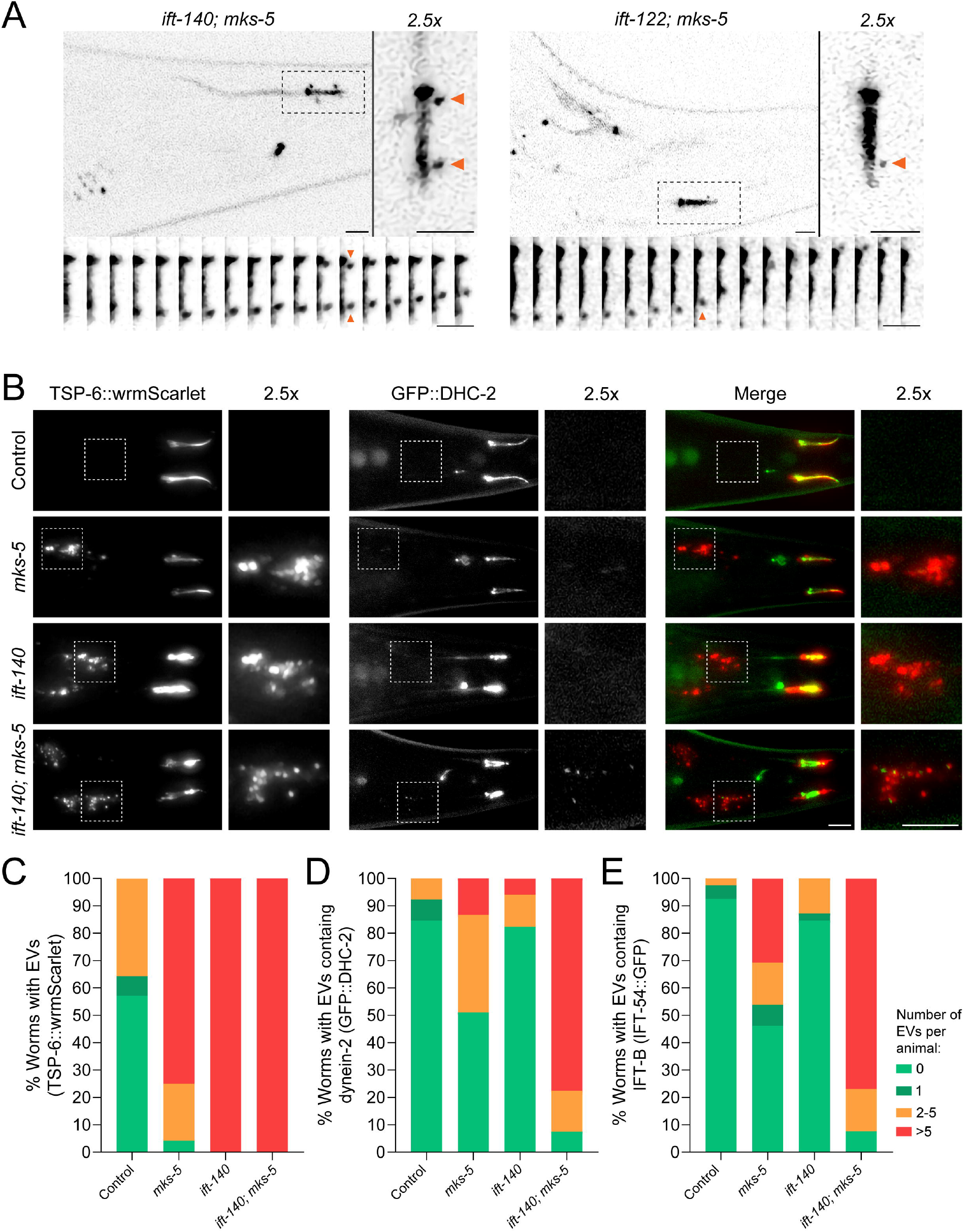
Loss of the TZ triggers ectosome shedding of IFT machinery along IFT-A-deficient cilia. **(A)** Time-lapse imaging of phasmid cilia expressing GFP::DHC-2 in control, IFT-A, *mks-5*, single and double mutant animals. Arrows indicate GFP::DHC-2-positive vesicle-like structures shedding from different ciliary locations. **(B)** Tails of animals co-expressing the ciliary membrane and ectosome maker TSP-6::wrmScarlet with GFP::DHC-2. Insets highlight ectosome accumulation in *mks-5* and IFT-A single and double mutants, with colocalization of GFP::DHC-2 observed in double mutants. **(C-E)** Quantification of animals of the indicated genotypes presenting ciliary EVs released from their phasmid cilia labelled by TSP-6::wrmScarlet (C), GFP::DHC-2 (D), or IFT-54::GFP (E). Percentage of animals without detectable EVs is shown in light green. Percentages of animals containing 1 ciliary EV (typically at the tip), 2-5 ciliary EVs (orange), or more than 5 ciliary EVs (often agglomerated in neighboring glia cells) are shown in dark green, orange, and red, respectively. (n≥39 animals per condition tested). Scale bars, 5 μm.

*C. elegans* phasmid cilia have been shown to release ciliary EVs from tips and the periciliary membrane compartment, which are taken up by neighboring glia, providing a tractable in vivo clearance sink (Akella et al., 2020; Lobo et al., 2025; Razzauti and Laurent, 2021; Wang et al., 2024; Wang et al., 2014). To determine whether the vesicle-like structures carrying DHC-2/IFT-54 that we observed corresponded to ciliary EVs, we examined the ciliary membrane and EV marker TSP-6::wrmScarlet (the *C. elegans* ortholog of CD9)(Razzauti and Laurent, 2021)(Figure 4B).

Control animals had none or a very small number of detectable TSP⍰6-positive EVs (1-5), which in most cases lacked IFT machinery (Figure 4C). In contrast, *mks-5* and *ift-140* single and double mutants exhibited an increase in TSP⍰6-labelled vesicles in the animals’ lumen: 75% of *mks-5* and 100% of *ift-140* single mutant animals had 6 or more TSP-6 positive ciliary EVs, and double mutants consistently displayed at least 6 of these vesicles (Figure 4C).

Importantly, while ∼50% of *mks-5* and 80% of *ift-140* single mutant animals had no DHC-2 or IFT-54 particles detectable in their ciliary EVs, these IFT subunits were readily detectable inside EVs in 95% of *mks-5*;*ift-140* double mutant animals (Figures 4D and 4E). Nonetheless, not all TSP-6–positive ciliary EVs contained DHC-2, indicating heterogeneity in ectosome composition in these double mutant animals (Figure 4B, last panel).

Together, these findings demonstrate that combined loss of IFT-A and the TZ not only promotes the release of ciliary EVs but also drives their abnormal enrichment with IFT machinery. This provides an additional route to relieve ciliary protein overload in the absence of the TZ to support axoneme extension when retrograde transport is compromised.

## DISCUSSION

Our results show that complete disruption of the ciliary TZ, via *mks-5* deletion, can rescue key defects of IFT-A-deficient cilia. In *ift-140* and *ift-122* mutants, where the IFT-A complex is lost and retrograde IFT is essentially abolished (Goncalves-Santos et al., 2023), TZ removal not only clears IFT accumulations but also improves axoneme extension. This rescue occurs without reinstating dynein-2–mediated retrograde transport; instead, clearance of IFT machinery is achieved through passive diffusion and release in ciliary EVs. These alternative disposal routes ultimately facilitate ciliogenesis.

### MKS-dependent TZ architecture blocks the passive exit of inactive dynein-2

Our findings corroborate previous studies that the TZ barrier controls not only entry but also the exit of IFT particles (De-Castro et al., 2022; Jensen et al., 2015; Prevo et al., 2015). We previously established that the TZ’s NPHP module poses a mechanical barrier to “underpowered” retrograde IFT trains when dynein-2 availability is reduced by loss of its intermediate chain, WDR-60 (De-Castro et al., 2022). Here, we find that NPHP-module disruption alone is insufficient to clear accumulated IFT machinery from cilia of retrograde IFT-deficient IFT-A mutants. Instead, IFT train clearance was possible only by completely blocking TZ assembly through the deletion of *mks-5*. Thus, our results suggest that even the MKS module by itself is sufficient to confer a barrier to IFT assemblies and inactive motors, impeding their passive ciliary exit in the absence of retrograde transport.

### Diffusion-driven IFT clearance as a compensatory mechanism in IFT-A/TZ double mutants

Consistent with our prior study (Goncalves-Santos et al., 2023), we observed increased anterograde IFT kinetics associated with the loss of the IFT-A complex. This effect may arise from the reduction in the size of anterograde IFT trains, as they no longer carry IFT-A nor components normally linked to this complex, such as the BBSome and TULP adaptors (Mukhopadhyay et al., 2010; Wei et al., 2012). Disruption of MKS-5 also increases proximal anterograde velocity, consistent with the loss of the TZ reducing mechanical resistance to IFT train passage, allowing them to enter the axoneme and reach maximum velocity at a faster rate (De-Castro et al., 2022; Jensen et al., 2015; Prevo et al., 2015).

Importantly, single-molecule imaging revealed that, in the absence of IFT-A, retrograde-bound IFT particles lose processivity and exhibit stochastic, irregular, zig-zag trajectories. These are consistent with diffusion-dominated behavior rather than motor-directed retrograde transport. In both *ift-140* and *ift-122* mutants, the non-processive particles remain trapped within the ciliary lumen; by contrast, when these mutants are combined with the *mks-5* mutant background, the diffusing IFT particles are able to exit cilia and no longer accumulate inside. This indicates that, under conditions where dynein-2-dependent motility is abolished, diffusion through a compromised TZ mitigates IFT particle accumulation inside cilia. We speculate that the continuous influx of new anterograde trains into an already overcrowded cilium may also promote the exit of inactive IFT assemblies once the TZ barrier is lost in the IFT-A mutant backgrounds. This possibility is supported by prior studies collectively showing that inward IFT train flow actively shapes diffusion rates and concentration gradients in cilia, implying that active anterograde transport can also favor outward diffusion of IFT components through physical and concentration effects (Patel et al., 2024; van Krugten et al., 2022; Zhang et al., 2021).

Collectively, these results indicate that, in the absence of both IFT-A and TZ, passive diffusion becomes a key mechanism for IFT particle clearance from cilia, possibly augmented by anterograde flow.

### Ciliary EV shedding as a complementary disposal pathway

In parallel with diffusion-mediated exit, we find that accumulated IFT machinery is abnormally incorporated and released inside ciliary EVs in IFT-A/TZ double mutants. Although single mutants also show elevated EV release (also shown recently in (Lobo et al., 2025)), only the double mutants display an enrichment of DHC-2 and IFT-54 within EVs, indicating that TZ integrity is a key determinant of ciliary EV cargo composition when retrograde IFT is lost.

While ciliary EV release is a conserved, regulated process in many cilia types, its cargo is usually biased toward receptors and signaling factors (Phua et al., 2017; Wang et al., 2024). The striking enrichment of IFT components in EVs from IFT-A/TZ double mutants therefore implies a stress-induced remodeling of cargo selection, allowing the cilium to export otherwise retained IFT complexes. This could be a failsafe mechanism to package and discard bulky, traffic jam-causing IFT assemblies when both retrograde transport and the TZ barrier are disrupted. Work in *Chlamydomonas* has also shown that loss of TZ components can alter ciliary EV/ectosome biogenesis and cargo sorting (Wang et al., 2022), supporting a conserved role for the TZ gate in coordinating membrane-budding events.

Whether EV-mediated disposal of IFT machinery is actively signaled (e.g. via post-translational modification of cargo or recruitment of membrane-remodeling factors) or is a passive consequence of altered ciliary membrane dynamics remains unclear. Nonetheless, our data show that TZ loss enables two complementary clearance mechanisms for accumulated IFT machinery: diffusion and ciliary EV-mediated export.

### Ciliary axoneme elongation without TZ and retrograde IFT

Mechanistically, we propose that diffusion- and EV-mediated export of IFT particles together enable the increased axoneme elongation observed in IFT-A/TZ double mutants. By clearing accumulated IFT assemblies, these mechanisms reduce crowding in the ciliary lumen. This likely facilitates the continued delivery of structural components, such as tubulin, and their incorporation at the ciliary tip even when canonical dynein-2/IFT-A– mediated retrograde transport is non-functional. The incomplete rescue of axoneme elongation in IFT-A/TZ double mutants suggests that the loss of ciliary compartmentalization and axoneme-membrane coupling may nonetheless reduce the efficiency of axoneme extension compared to wild-type cilia (Jensen et al., 2015).

Given the wide conservation of IFT and TZ modules, as well as the broad presence of ciliary EVs from *C. elegans* to vertebrates, the mechanisms identified here likely represent generalizable principles applicable to diverse cilia and flagella. Interestingly, however, evolutionary exceptions exist in which retrograde transport and canonical TZ are not required for axoneme assembly. For instance, *Drosophila* sperm assemble axonemes within the cytoplasm and develop a membrane-associated TZ “cap” only during later stages of flagellar formation, effectively bypassing requirement for retrograde IFT (Basiri et al., 2014; Briggs et al., 2004). Consistent with this, mutations in IFT components compromise sensory cilia but not sperm flagella in *Drosophila* (Han et al., 2003; Sarpal et al., 2003), highlighting divergent dependencies on IFT and TZ architecture in different cell types (Akella et al., 2019; Jana et al., 2018). A more extreme example is *Plasmodium* species, which lack canonical IFT machinery entirely and assemble their gamete flagellar axonemes in the cytoplasm before plasma membrane docking, without evident TZ structures (Briggs et al., 2004; Sinden et al., 2010). Our data suggest that *C. elegans* cilia can similarly adapt to the absence of both TZ and retrograde IFT, activating unconventional disposal routes to sustain cilia elongation.

Future work should assess whether selective, partial modulation of TZ permeability can be harnessed therapeutically to treat ciliopathies associated with retrograde IFT defects without entirely compromising TZ function and ciliary signaling.

## Limitations

This study was performed in *C. elegans* sensory cilia, an animal model notable for genetic tractability and vivo imaging of IFT. While the outlined ciliary mechanisms are broadly conserved, systematic studies across species will be necessary to fully elucidate the universality and physiological impact of these findings.

## ACKNOWLEDGEMENTS

We thank Dr Ana Carvalho for sharing equipment and for exchanging ideas. We are also very grateful to Dr Guangshuo Ou, Dr Patrick Laurent and Dr Jessica Feldman for providing *C. elegans* strains.

This work was supported by the projects 2023.12458.PEX (to C.M.C.A.) and 2022.01955.PTDC (to T.J.D.) from the Fundação para a Ciência e a Tecnologia (FCT). This work has also been funded by national funds, through the FCT, under projects UID/06304/2023 and LA/P/0050/2020 (DOI 10.54499/LA/P/0050/2020).

A.R.G.D.-C. and M.J.G.D-C received PhD fellowships from FCT (UI/BD/152865/2022 and 2023.01378.BD, respectively) and support from the Molecular and Cell Biology, and the Animal Sciences PhD programs at ICBAS.

R.G, C.M.C.A., and T.J.D. salaries were also supported by FCT: CEECIND/00333/2017, CEECIND/01985/2018 and CEECINSTLA/00012/2022,respectively.

E.J.G.P. and G.H. acknowledge support by the Dutch Research Council (NWO; Project no. OCENW.M20.063).

The authors also thank the National Bioresource Project for *C. elegans* and the *Caenorhabditis* Genetics Center (CGC) for providing strains.

## AUTHOR CONTRIBUTIONS

A.R.G.D. performed most of the experiments, actively contributed to the experimental design, analyzed data, prepared figures and wrote the original draft of the manuscript. M.J.G.D. helped with some experiments and figure preparation. G.H.H. helped with single-molecule live imaging experiments. R.G. and E.J.G.P. provided strains, reagents, equipment, and helped interpreting results. T.J.D. and C.M.C.A. conceived and supervised the project, designed and helped with experiments, analyzed and interpreted data, and helped to prepare figures and to write the manuscript. All authors read the manuscript and provided input for the final version.

## DECLARATION OF INTERESTS

The authors declare no competing interests.

## FIGURE LEGENDS

**Figure S1. NPHP-4 disruption fails to clear IFT accumulations in cilia from IFT-A mutants**

**(A)** Phasmid cilia from control, IFT-A1, *nphp-4*, single and double mutant animals expressing GFP::DHC-2. Scale bar: 2 μm.

**(B-C)** Distribution profiles of GFP::DHC-2 along cilia of the indicated genotypes (n≥38 cilia per condition tested). For IFT-A single and IFT-A;*nphp-4* double mutants, average intensity profiles are shown for two observed ciliary phenotype categories, as well as the combined average profile of all cilia. The ciliary base is set at position 0 (dotted gray line). Data represent show mean ± SEM.

**Figure S2. Anterograde IFT reaches maximum speed earlier in IFT-A-deficient cilia**

**(A, B)** Anterograde velocity of GFP::DHC-2 particles at different positions along cilia in control versus IFT-A single mutant animals (A; n ≥ 50 particle traces per cilium in ≥ 48 animals, per condition tested), and *mks-5* single mutant versus IFT-A;*mks-5* double mutants (B; n ≥ 50 particle traces per cilium in ≥ 34 animals, per condition tested). Graphs show mean ± SEM.

**Figure S3. TZ removal enables passive diffusion of stationary IFT trains in retrograde-deficient mutants**

**(A)** Kymographs of endogenous GFP::DHC-2, in control, IFT-A, *mks-5*, single and double mutant backgrounds following sequential photobleaching. IFT particles form a diffuse fluorescent “cloud” after reaching the ciliary tip in IFT-A;*mks-5* double mutants. The first and second photobleaching events indicated in the figure correspond to the bleaching of the signal inside the cilium body and the ciliary base, respectively. Scale bars: vertical, 5 s; horizontal, 2 μm.

**(B)** Kymographs of IFT-54::GFP in control, IFT-A, *mks-5*, single and double mutant backgrounds obtained upon single particle fluorescence imaging. Anterograde and retrograde particle trajectories are shown in red and green, respectively. Trajectories exhibiting diffusion-like motion are shown in orange. Scale bars: vertical, 2s; horizontal, 2 μm.

## MATERIALS AND METHODS

### Maintenance and generation of *Caenorhabditis elegans* strains

*Caenorhabditis elegans* strains were maintained under standard laboratory conditions at 20°C on Nematode Growth Medium (NGM) agar plates seeded with Escherichia coli OP50 as a food source. To maintain optimal growth and avoid overcrowding, adult hermaphrodites were transferred to fresh plates every 3–4 days. Due to their ability to self-fertilize, *C. elegans* hermaphrodites allowed for the propagation of genetically stable homozygous lines without the need for continuous mating. For long-term storage, strains were cryopreserved in 15% glycerol and stored at −80°C.

To combine already available mutations, fluorescent markers, and transgenes into genetic backgrounds of interest, classical genetic crossing was used by mating young adult males with hermaphrodites following standard protocols, and the progeny were selected through PCR-based genotyping. The strains and primers used are listed in the key resources table.

### *C. elegans* genome editing

For introducing new mutations or fluorescent tags, genome engineering tools such as CRISPR-Cas9 or MosSCI with native promoters were employed. For CRISPR-based edits, guide RNAs (gRNAs) were designed using the “CRISPOR” platform to target specific loci. gRNAs, Cas9 protein along with homology repair templates were microinjected into the gonads of young adult hermaphrodites. Homology repair templates were constructed as single-stranded oligonucleotides or plasmid-based donors containing homology arms (∼1000 bps) flanking the desired insertion site. Successfully edited strains were validated via PCR and sequencing. In order to remove any unintended mutations (off-targets), all edited strains were outcrossed to the wild-type N2 strain for at least four to five generations. All strains, primers, and guide RNAs used are detailed in the key resources table.

### Fluorescence imaging of *C. elegans* cilia

All imaging was performed using phasmid cilia of young adult *C. elegans* hermaphrodites expressing fluorescently tagged IFT components from knock-ins or single-copy insertion (MosSCI) transgenes with native promoters. Prior to imaging, animals were immobilized using 10 mM levamisole and mounted on 5% agarose pads on standard glass slides. All experiments were conducted in temperature-controlled rooms maintained at 20□°C to preserve physiological conditions.

### Ciliary measurements and quantifications

For quantitation of ciliary signal intensity profiles, high-resolution imaging was performed using an Axio Observer epifluorescence microscope (Zeiss). The microscope was equipped with a Plan-Apochromat 63×/1.46 NA oil immersion objective and an Orca Flash 4.0 camera (Hamamatsu). Image acquisition was done using ZEN software (Zeiss), and Z-stacks were acquired with 0.4 µm intervals.

Acquired z-stack and time-lapse datasets were processed using Fiji software (ImageJ v2.1.0/1.52v). The distribution profiles of IFT components were quantified by drawing a line along the length of each cilium, from the base to the distal tip, and then by extracting pixel intensity values along the drawn line. To correct for non-specific fluorescence, a background area near each cilium was measured and its signal subtracted from the raw data. Intensity values from numerous cilia were subsequently averaged and plotted as a function of distance from the base. In addition, the signal was normalized using a background area that was manually drawn outside the worm. This region was chosen to represent a baseline level, free from any biological signal, ensuring that the normalization process accounted only for background intensity. This approach helps to reduce variations in light intensity over time and noise and variability across samples, without altering or distorting the actual biological signal from the worm.

Ciliary length was quantitatively measured in both wild-type and mutant strains using fluorescently tagged proteins, such as GFP::DHC-2 and TBB-4::GFP. For each cilium, a line was drawn from the base of the cilium to its tip to determine its length.

The number of ciliary EVs was quantified manually using three different fluorescent markers: TSP-6::wrmScarlet, DHC-2::GFP, and IFT-54::GFP. For each animal analyzed, EVs were scored based on their visible fluorescence signal. To facilitate comparison across samples, cilia were categorized into four groups according to the number of EVs detected: no EVs(0), one EV (1; normally at the tip), two to five EVs (2-5), and more than five EVs (>5).

### Live imaging quantifications

Dynamic IFT was imaged using an Olympus IX81 inverted microscope integrated with an Andor Revolution XD spinning disk confocal system. This setup incorporated a CSU-X1 scanner (Yokogawa Electric Corp.), a solid-state laser combiner (ALC-UVP 350i; Andor Technology), and an iXonEM+ DU-897 EMCCD camera with a 2× port coupler, which were all controlled by Andor iQ3 software (Andor Technology). Time-lapse sequences were acquired at 3 frames per second (333 ms per frame) and consisted of 200 frames per phasmid cilium using a UPLSAPO 100×/1.40 NA oil objective.

To study IFT dynamics, anterograde and retrograde IFT kymographs were generated using the KymographClear plugin (version 2.0; (Mangeol et al., 2016)) in ImageJ. Only cilia completely focused in a single focal plane and without any strong morphological defects were taken into consideration to guarantee the quality of the data used to study IFT dynamics. Kymograph analysis was performed using KymographDirect (version 2.1; (Mangeol et al., 2016)), which corrects for photobleaching and background noise, as well as accurately separating anterograde from retrograde transport events. IFT particle tracks were automatically detected and validated manually to ensure the correct quantification as in (De-Castro et al., 2022; Goncalves-Santos et al., 2023). KymographDirect was also used to calculate particle intensities along the cilium and their average velocity.

### Sequential Photobleaching

Photobleaching was conducted on the Andor Revolution XD spinning disk confocal system using the 405 nm laser (60 mW) at 70% power in combination with a 60x/1.40 NA oil-immersion Plan-Apochromat objective. The initial photobleaching event targeted the axonemal region of the cilium to eliminate all GFP::DHC-2 fluorescence along its length. After a short delay (∼3 seconds) to allow IFT particles to enter the cilium, a second bleaching event was performed at the ciliary base. This two-step approach enabled the tracking of a defined number of particles over time, including in cilia with high levels of IFT accumulation. This photobleaching and imaging approach is described in detail in (Goncalves-Santos et al., 2023), which provides a comprehensive explanation of the technique and its applications.

### Single-Particle Live Imaging

For single molecule analyses at high temporal resolution, a Nikon Ti-E inverted microscope was used, fitted with a high numerical aperture 100x oil immersion objective(Nikon CFI Apo TIRF 100x/ NA 1.49). System operation and image acquisition were managed using MicroManager software (version 1.4). Fluorescence excitation was achieved with a 491 nm DPSS laser (Cobolt Calypso, 50 mW), and laser intensity was modulated through an acousto-optic tunable filter (AOTF; AA Optoelectronics). The diaphragm opening was manually set to regulate illumination area. Separation of excitation and emission light was accomplished using a dichroic mirror (ZT 405/488/561rpc, Chroma) in combination with a 525/45 nm bandpass emission filter (Brightline HC, Semrock). Image capture was performed using an EMCCD camera (iXon 897; Andor Technology).

For single-molecule imaging, a 10–15 μm region centered on a ciliary pair was selectively illuminated. Continuous exposure to high-intensity laser light led to the photobleaching of the majority of fluorescent molecules, thereby isolating a specific population for single-molecule resolution as described in (Mitra et al., 2024). Once this was achieved, the laser power was reduced to maintain the single-molecule state without prematurely bleaching the remaining fluorophores.

### Statistical analysis

Statistical analyses were conducted using GraphPad Prism (version 9). Each dataset was first assessed for normality using a range of tests, including the Anderson-Darling, D’Agostino-Pearson, Shapiro-Wilk, and Kolmogorov-Smirnov tests, to determine whether the data followed a Gaussian distribution. This guided the decision between applying parametric or non-parametric statistical tests. For comparisons between two groups, either the two-tailed Student’s t test (for normally distributed data) or the Mann-Whitney U test (for non-normal data) was used. When comparing more than two groups, one-way ANOVA was applied for parametric data and the Kruskal-Wallis test for non-parametric data, followed by appropriate post hoc multiple comparisons. Statistical significance was defined as p ≤ 0.05, with the following notations: p ≤ 0.05 (*), p ≤ 0.01 (**), p ≤ 0.001 (***), and p ≤ 0.0001 (****). Graphs representing XY position-averaged velocity and intensity distributions display data as mean ± SEM, while bar charts show mean ± SD.

## Notes

### Competing Interest Statement

The authors have declared no competing interest.

## REFERENCES

Akella, J. S., Carter, S. P., Nguyen, K., Tsiropoulou, S., Moran, A. L., Silva, M., Rizvi, F., Kennedy, B. N., Hall, D. H., Barr, M. M. et al. (2020). Ciliary Rab28 and the BBSome negatively regulate extracellular vesicle shedding. Elife 9.

Akella, J. S., Silva, M., Morsci, N. S., Nguyen, K. C., Rice, W. J., Hall, D. H. and Barr, M. M. (2019). Cell type-specific structural plasticity of the ciliary transition zone in C. elegans. Biol Cell 111, 95–107.

Basiri, M. L., Ha, A., Chadha, A., Clark, N. M., Polyanovsky, A., Cook, B. and Avidor-Reiss, T. (2014). A migrating ciliary gate compartmentalizes the site of axoneme assembly in Drosophila spermatids. Curr Biol 24, 2622–31.

Briggs, L. J., Davidge, J. A., Wickstead, B., Ginger, M. L. and Gull, K. (2004). More than one way to build a flagellum: comparative genomics of parasitic protozoa. Curr Biol 14, R611–2.

De-Castro, A. R. G., Rodrigues, D. R. M., De-Castro, M. J. G., Vieira, N., Vieira, C., Carvalho, A. X., Gassmann, R., Abreu, C. M. C. and Dantas, T. J. (2022). WDR60-mediated dynein-2 loading into cilia powers retrograde IFT and transition zone crossing. J Cell Biol 221.

Garcia-Gonzalo, F. R. and Reiter, J. F. (2017). Open Sesame: How Transition Fibers and the Transition Zone Control Ciliary Composition. Cold Spring Harb Perspect Biol 9.

Goncalves-Santos, F., De-Castro, A. R. G., Rodrigues, D. R. M., De-Castro, M. J. G., Gassmann, R., Abreu, C. M. C. and Dantas, T. J. (2023). Hot-wiring dynein-2 establishes roles for IFT-A in retrograde train assembly and motility. Cell Rep 42, 113337.

Han, Y. G., Kwok, B. H. and Kernan, M. J. (2003). Intraflagellar transport is required in Drosophila to differentiate sensory cilia but not sperm. Curr Biol 13, 1679–86.

Hao, L., Thein, M., Brust-Mascher, I., Civelekoglu-Scholey, G., Lu, Y., Acar, S., Prevo, B., Shaham, S. and Scholey, J. M. (2011). Intraflagellar transport delivers tubulin isotypes to sensory cilium middle and distal segments. Nat Cell Biol 13, 790–8.

Hesketh, S. J., Mukhopadhyay, A. G., Nakamura, D., Toropova, K. and Roberts, A. J. (2022). IFT-A structure reveals carriages for membrane protein transport into cilia. Cell 185, 4971–4985 e16.

Jana, S. C., Mendonca, S., Machado, P., Werner, S., Rocha, J., Pereira, A., Maiato, H. and Bettencourt-Dias, M. (2018). Differential regulation of transition zone and centriole proteins contributes to ciliary base diversity. Nat Cell Biol 20, 928–941.

Jensen, V. L., Lambacher, N. J., Li, C., Mohan, S., Williams, C. L., Inglis, P. N., Yoder, B. K., Blacque, O. E. and Leroux, M. R. (2018). Role for intraflagellar transport in building a functional transition zone. EMBO Rep 19.

Jensen, V. L., Li, C., Bowie, R. V., Clarke, L., Mohan, S., Blacque, O. E. and Leroux, M. R. (2015). Formation of the transition zone by Mks5/Rpgrip1L establishes a ciliary zone of exclusion (CIZE) that compartmentalises ciliary signalling proteins and controls PIP2 ciliary abundance. EMBO J 34, 2537–56.

Kozminski, K. G., Johnson, K. A., Forscher, P. and Rosenbaum, J. L. (1993). A motility in the eukaryotic flagellum unrelated to flagellar beating. Proc Natl Acad Sci U S A 90, 5519–23.

Lacey, S. E. and Pigino, G. (2025). The intraflagellar transport cycle. Nat Rev Mol Cell Biol 26, 175–192.

Lobo, T., Haasnoot, G. H., Nawrocka, A., Bruggeman, C. W., Razzauti, A., Peterman, E. J. G. and Laurent, P. (2025). Sensory stimuli and cilium trafficking defects trigger the release of ciliary extracellular vesicles from multiple ciliary locations. bioRxiv.

Magescas, J., Eskinazi, S., Tran, M. V. and Feldman, J. L. (2021). Centriole-less pericentriolar material serves as a microtubule organizing center at the base of C. elegans sensory cilia. Curr Biol 31, 2410–2417 e6.

Mangeol, P., Prevo, B. and Peterman, E. J. (2016). KymographClear and KymographDirect: two tools for the automated quantitative analysis of molecular and cellular dynamics using kymographs. Mol Biol Cell 27, 1948–57.

Mill, P., Christensen, S. T. and Pedersen, L. B. (2023). Primary cilia as dynamic and diverse signalling hubs in development and disease. Nat Rev Genet 24, 421–441.

Mitra, A., Loseva, E. and Peterman, E. J. G. (2024). IFT cargo and motors associate sequentially with IFT trains to enter cilia of C. elegans. Nat Commun 15, 3456.

Moran, A. L., Louzao-Martinez, L., Norris, D. P., Peters, D. J. M. and Blacque, O. E. (2024). Transport and barrier mechanisms that regulate ciliary compartmentalization and ciliopathies. Nat Rev Nephrol 20, 83–100.

Mukhopadhyay, S., Wen, X., Chih, B., Nelson, C. D., Lane, W. S., Scales, S. J. and Jackson, P. K. (2010). TULP3 bridges the IFT-A complex and membrane phosphoinositides to promote trafficking of G protein-coupled receptors into primary cilia. Genes Dev 24, 2180–93.

Nager, A. R., Goldstein, J. S., Herranz-Perez, V., Portran, D., Ye, F., Garcia-Verdugo, J. M. and Nachury, M. V. (2017). An Actin Network Dispatches Ciliary GPCRs into Extracellular Vesicles to Modulate Signaling. Cell 168, 252–263 e14.

Nakamura, K., Noguchi, T., Takahara, M., Omori, Y., Furukawa, T., Katoh, Y. and Nakayama, K. (2020). Anterograde trafficking of ciliary MAP kinase-like ICK/CILK1 by the intraflagellar transport machinery is required for intraciliary retrograde protein trafficking. J Biol Chem 295, 13363–13376.

Nechipurenko, I. and Sengupta, P. (2025). C. elegans: An elegant experimental system for the study of cilia biology. Semin Cell Dev Biol 174, 103636.

Patel, M. B., Griffin, P. J., Olson, S. F., Dai, J., Hou, Y., Malik, T., Das, P., Zhang, G., Zhao, W., Witman, G. B. et al. (2024). Distribution and bulk flow analyses of the intraflagellar transport(IFT) motor kinesin-2 support an “on-demand” model for Chlamydomonas ciliary length control. Cytoskeleton (Hoboken) 81, 586–604.

Pazour, G. J., Dickert, B. L. and Witman, G. B. (1999). The DHC1b (DHC2) isoform of cytoplasmic dynein is required for flagellar assembly. J Cell Biol 144, 473–81.

Phua, S. C., Chiba, S., Suzuki, M., Su, E., Roberson, E. C., Pusapati, G. V., Schurmans, S., Setou, M., Rohatgi, R., Reiter, J. F. et al. (2017). Dynamic Remodeling of Membrane Composition Drives Cell Cycle through Primary Cilia Excision. Cell 168, 264–279 e15.

Picariello, T., Brown, J. M., Hou, Y., Swank, G., Cochran, D. A., King, O. D., Lechtreck, K., Pazour, G. J. and Witman, G. B. (2019). A global analysis of IFT-A function reveals specialization for transport of membrane-associated proteins into cilia. J Cell Sci 132.

Piperno, G., Siuda, E., Henderson, S., Segil, M., Vaananen, H. and Sassaroli, M. (1998). Distinct mutants of retrograde intraflagellar transport (IFT) share similar morphological and molecular defects. J Cell Biol 143, 1591–601.

Prevo, B., Mangeol, P., Oswald, F., Scholey, J. M. and Peterman, E. J. (2015). Functional differentiation of cooperating kinesin-2 motors orchestrates cargo import and transport in C. elegans cilia. Nat Cell Biol 17, 1536–45.

Razzauti, A. and Laurent, P. (2021). Ectocytosis prevents accumulation of ciliary cargo in C. elegans sensory neurons. Elife 10.

Reddy Palicharla, V. and Mukhopadhyay, S. (2024). Molecular and structural perspectives on protein trafficking to the primary cilium membrane. Biochem Soc Trans 52, 1473–1487.

Reiter, J. F. and Leroux, M. R. (2017). Genes and molecular pathways underpinning ciliopathies. Nat Rev Mol Cell Biol 18, 533–547.

Salinas, R. Y., Pearring, J. N., Ding, J. D., Spencer, W. J., Hao, Y. and Arshavsky, V. Y. (2017). Photoreceptor discs form through peripherin-dependent suppression of ciliary ectosome release. J Cell Biol 216, 1489–1499.

Sarpal, R., Todi, S. V., Sivan-Loukianova, E., Shirolikar, S., Subramanian, N., Raff, E. C., Erickson, J. W., Ray, K. and Eberl, D. F. (2003). Drosophila KAP interacts with the kinesin II motor subunit KLP64D to assemble chordotonal sensory cilia, but not sperm tails. Curr Biol 13, 1687–96.

Scheidel, N. and Blacque, O. E. (2018). Intraflagellar Transport Complex A Genes Differentially Regulate Cilium Formation and Transition Zone Gating. Curr Biol 28, 3279–3287 e2.

Sinden, R. E., Talman, A., Marques, S. R., Wass, M. N. and Sternberg, M. J. (2010). The flagellum in malarial parasites. Curr Opin Microbiol 13, 491–500.

van Krugten, J., Danne, N. and Peterman, E. J. G. (2022). A local interplay between diffusion and intraflagellar transport distributes TRPV-channel OCR-2 along C. elegans chemosensory cilia. Commun Biol 5, 720.

Wang, J., Barr, M. M. and Wehman, A. M. (2024). Extracellular vesicles. Genetics 227.

Wang, J., Silva, M., Haas, L. A., Morsci, N. S., Nguyen, K. C., Hall, D. H. and Barr, M. M. (2014). C. elegans ciliated sensory neurons release extracellular vesicles that function in animal communication. Curr Biol 24, 519–25.

Wang, L., Wen, X., Wang, Z., Lin, Z., Li, C., Zhou, H., Yu, H., Li, Y., Cheng, Y., Chen, Y. et al. (2022). Ciliary transition zone proteins coordinate ciliary protein composition and ectosome shedding. Nat Commun 13, 3997.

Wei, Q., Zhang, Y., Li, Y., Zhang, Q., Ling, K. and Hu, J. (2012). The BBSome controls IFT assembly and turnaround in cilia. Nat Cell Biol 14, 950–7.

Williams, C. L., Li, C., Kida, K., Inglis, P. N., Mohan, S., Semenec, L., Bialas, N. J., Stupay, R. M., Chen, N., Blacque, O. E. et al. (2011). MKS and NPHP modules cooperate to establish basal body/transition zone membrane associations and ciliary gate function during ciliogenesis. J Cell Biol 192, 1023–41.

Wood, C. R., Huang, K., Diener, D. R. and Rosenbaum, J. L. (2013). The cilium secretes bioactive ectosomes. Curr Biol 23, 906–11.

Zhang, Z., Danne, N., Meddens, B., Heller, I. and Peterman, E. J. G. (2021). Direct imaging of intraflagellar-transport turnarounds reveals that motors detach, diffuse, and reattach to opposite-direction trains. Proc Natl Acad Sci U S A 118.

