## Supplementary material for "Removal of the Ciliary Gate Allows Axoneme Extension in the Absence of Retrograde IFT": Figures S1 to S3 + Tables

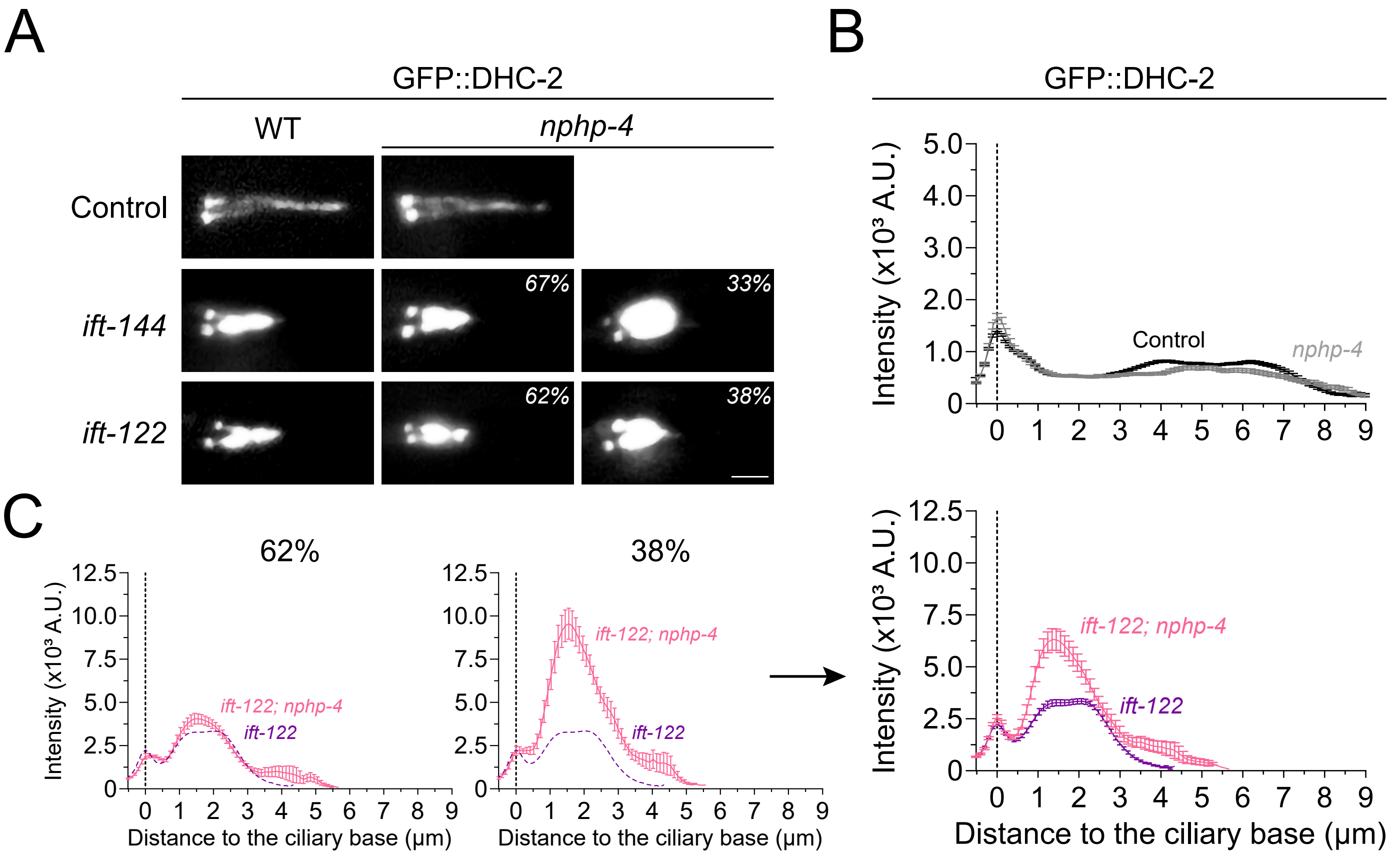


**Figure S1. NPHP-4 disruption fails to clear IFT accumulations in cilia from IFT-A mutants.**
**(A)** Phasmid cilia from control, IFT-A, *nphp-4* single and double mutant animals expressing GFP::DHC-2. Scale bar: 2 μm. **(B-C)** Distribution profiles of GFP::DHC-2 along cilia of the indicated genotypes (n≥38 cilia per condition tested). For IFT-A and IFT-A; *nphp-4* double mutants, average intensity profiles are shown for two observed ciliary phenotype categories, as well as the combined average profile of all cilia. The ciliary base is set at position 0 (dotted gray line). Data represent show mean ± SEM.

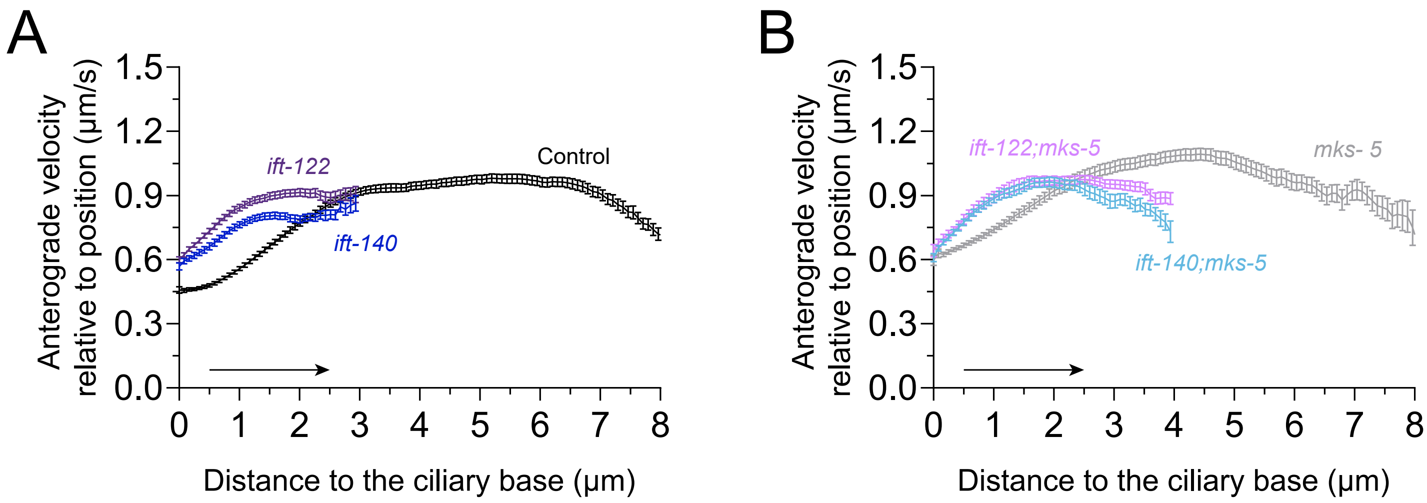


**Figure S2. Anterograde IFT reaches maximum speed earlier in IFT-A-deficient cilia.**

**(A, B)** Anterograde velocity of GFP::DHC-2 particles at different positions along cilia in control versus IFT-A1 single mutant animals (A; n ≥ 50 particle traces per cilium in ≥ 48 animals per condition tested), and *mks-5* single mutant versus IFT-A1;*mks-5* double mutant (B; n ≥ 50 particle traces per cilium in ≥ 34 animals per condition tested). Graphs show mean ± SEM**.**

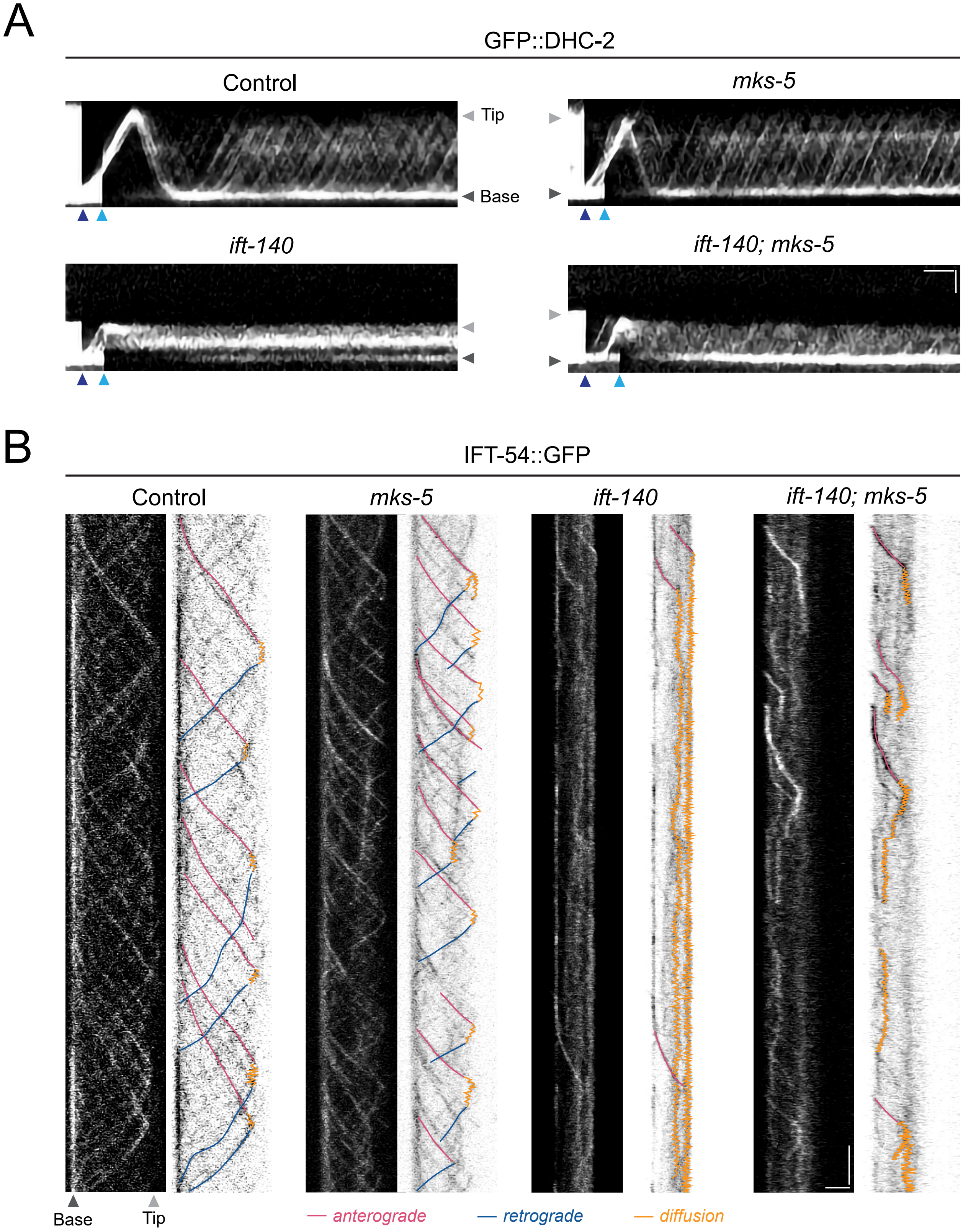


**Figure S3. TZ removal enables passive diffusion of stationary IFT trains in retrograde-deficient mutants.** **(A)** Kymographs of endogenous GFP::DHC-2, in control, IFT-A, *mks-5* single and double mutant backgrounds following sequential photobleaching. IFT particles form a diffuse fluorescent "cloud" after reaching the ciliary tip in IFT-A/TZ double mutants. The first and second photobleaching events indicated in the figure correspond to the bleaching of the signal inside the cilium body and the ciliary base, respectively. Scale bars: horizontal, 5 s; vertical, 2 µm. **(B)** Kymographs of IFT-54::GFP in control, IFT-A, *mks-5* single and double mutant backgrounds obtained upon single particle fluorescence imaging. Anterograde and retrograde particle trajectories are shown in magenta and blue, respectively. Trajectories exhibiting diffusion-like motion are shown in orange. Scale bars: vertical, 2s; horizontal, 2 µm. See also Videos SX.

**Table S1.** List of strains used in this study.

| Strains | Genotype | Short annotation | Reference |
|---|---|---|---|
| N2 | | wild-type | |
| EJP401 | *vuaSi401 [pSA401; Ptbb-4::tbb-4::eGFP;cb-unc119(+)] I* | TBB-4::GFP | Mijalkovic et al., 2018 |
| GOU2362 | *ift-74 (cas499[ift-74::gfp]) II* | IFT-74::GFP | Yi et al., 2017 |
| JLF425 | *spd-5 (wow36[tagRFP-T::spd-5]); spd-2(wow60[spd-2::GFP::3xFlag]) I* | SPD-5::RFP + SPD-2::GFP | Magescas et al. 2019 |
| AND12 | *dhc-2(cas443[gfp::dhc-2]) I* | GFP::DHC-2 | De-Castro et al., 2021 |
| AND47 | *wdr-60(dan3[wdr-60::linker::3xFlag::gfp]) III* | WDR-60::GFP | De-Castro et al., 2021 |
| AND91 | *dhc-2(cas443[gfp::dhc-2]) I; ift-140(tm3433) V* | GFP::DHC-2 + *ift-140* | Gonçalves et al. 2023 |
| AND102 | *wdr-60(dan3[wdr-60::linker::3xFlag::gfp]) III; ift-140(tm3433) V* | WDR-60::GFP + *ift-140* | Gonçalves et al. 2023 |
| AND173 | *dhc-2(cas443[gfp::dhc-2])I; nphp-4(tm925)V* | GFP::DHC-2 + *nphp-4* | De-Castro et al., 2021 |
| AND182 | *wdr-60(dan3[wdr-60::linker::3xFlag::gfp]) III; mks-5(tm3100) II* | WDR-60::GFP + *mks-5* | *this study* |
| AND188 | *dhc-2(cas443[gfp::dhc-2]) I; mks-5(tm3100) II; ift-140(tm3433) V* | GFP::DHC-2 + *mks-5* + *ift-140* | *this study* |
| AND194 | *dhc-2(cas443[gfp::dhc-2]) I; mks-5(tm3100) II* | GFP::DHC-2 + *mks-5* | De-Castro et al., 2021 |
| AND241 | *dhc-2(cas443[gfp::dhc-2]) I; ift-144(gk678) III* | GFP::DHC-2 + *ift-144* | Gonçalves et al. 2023 |
| AND248 | *dhc-2(cas443[gfp::dhc-2]) I; mks-5(tm3100) II; ift-122(tm2878) IV* | GFP::DHC-2 + *mks-5* + *ift-122* | *this study* |
| AND249 | *dhc-2(cas443[gfp::dhc-2]) I; ift-122(tm2878) IV* | GFP::DHC-2 + *ift-122* | Gonçalves et al. 2023 |
| AND278 | *wdr-60(dan3[wdr-60::linker::3xFlag::gfp]) III; ift-140(tm3433) V; mks-5(tm3100) II* | WDR-60::GFP + *mks-5* + *ift-140* | *this study* |
| AND282 | *wdr-60(dan3[wdr-60::linker::3xFlag::gfp]) III; ift-122(tm2878) IV; mks-5(tm3100) II* | WDR-60::GFP + *mks-5* + *ift-122* | *this study* |
| AND283 | *wdr-60(dan3[wdr-60::linker::3xFlag::gfp]) III; ift-122(tm2878) IV* | WDR-60::GFP + *ift-122* | Gonçalves et al. 2023 |
| AND291 | *ift-74(cas499[ift-74::gfp]) II; mks-5(tm3100) II* | IFT-74::GFP + *mks-5* | *this study* |
| AND307 | *ift-74(cas499[ift-74::gfp]) II; mks-5(tm3100) II; ift-140(tm3433) V* | IFT-74::GFP + *mks-5* + *ift-140* | *this study* |
| AND308 | *ift-74(cas499[ift-74::gfp]) II; mks-5(tm3100) II; ift-122(tm2878) IV* | IFT-74::GFP + *mks-5* + *ift-122* | *this study* |
| AND311 | *ift-74(cas499[ift-74::gfp]) II; ift-140(tm3433) V* | IFT-74::GFP + *ift-140* | Gonçalves et al. 2023 |
| AND312 | *ift-74(cas499[ift-74::gfp]) II; ift-122(tm2878) IV* | IFT-74::GFP + *ift-122* | Gonçalves et al. 2023 |
| AND354 | *dhc-2(cas443[gfp::dhc-2])I; nphp-4(tm925) V; ift-144(gk678) III* | GFP::DHC-2 + *nphp-4* + *ift-144* | *this study* |
| AND355 | *dhc-2(cas443[gfp::dhc-2])I; nphp-4(tm925) V; ift-122(tm2878) IV* | GFP::DHC-2 + *nphp-4* + *ift-122* | *this study* |
| AND470 | *dan26 [ift-54::aid::gfp] X* | IFT-54::GFP | Gonçalves et al. 2023 |
| AND487 | *dan26 [ift-54::aid::gfp] X; ift-140(tm3433) V* | IFT-54::GFP + *ift-140* | Gonçalves et al. 2023 |
| AND489 | *dan26 [ift-54::aid::gfp] X; ift-122(tm2878) IV* | IFT-54::GFP + *ift-122* | Gonçalves et al. 2023 |
| AND567 | *dan26 [ift-54::aid::gfp] X; mks-5(tm3100) II* | IFT-54::GFP + *mks-5* | *this study* |
| AND568 | *dan26 [ift-54::aid::gfp] X; mks-5(tm3100) II; ift-140(tm3433) V* | IFT-54::GFP + *mks-5* + *ift-140* | *this study* |
| AND752 | *spd-5(wow36[tagRFP-T::spd-5]) I; vuaSi401 [pSA401; Ptbb-4::tbb-4::eGFP;cb-unc119(+)] I* | SPD-5::RFP + TBB-4::GFP | *this study* |
| AND760 | *tsp-6::wrmScarlet X; mks-5(tm3100) II; dhc-2(cas443[gfp::dhc-2]) I* | TSP-6::wrmScarlet + *mks-5* + GFP::DHC-2 | *this study* |
| AND761 | *tsp-6::wrmScarlet X; mks-5(tm3100) II; ift-140(tm3433) V; dhc-2(cas443[gfp::dhc-2]) I* | TSP-6::wrmScarlet + *mks-5* + *ift-140* + GFP::DHC-2 | *this study* |
| AND769 | *tsp-6::wrmScarlet X; dhc-2(cas443[gfp::dhc-2]) I* | TSP-6::wrmScarlet + GFP::DHC-2 | *this study* |
| AND770 | *tsp-6::wrmScarlet X; ift-140(tm3433) V; dhc-2(cas443[gfp::dhc-2]) I* | TSP-6::wrmScarlet + *ift-140* + GFP::DHC-2 | *this study* |
| AND771 | *spd-5(wow36[tagRFP-T::spd-5]) I; vuaSi401 [pSA401; Ptbb-4::tbb-4::eGFP;cb-unc119(+)] I; mks-5(tm3100) II* | SPD-5::RFP + TBB-4::GFP + *mks-5* | *this study* |
| AND772 | *spd-5(wow36[tagRFP-T::spd-5]) I; vuaSi401 [pSA401; Ptbb-4::tbb-4::eGFP;cb-unc119(+)] I; mks-5(tm3100) II; ift-140(tm3433) V* | SPD-5::RFP + TBB-4::GFP + *mks-5* + *ift-140* | *this study* |
| AND773 | *spd-5(wow36[tagRFP-T::spd-5]) I; vuaSi401 [pSA401; Ptbb-4::tbb-4::eGFP;cb-unc119(+)] I; ift-140(tm3433) V* | SPD-5::RFP + TBB-4::GFP + *ift-140* | *this study* |
| AND774 | *ift-54(dan26 [ift-54::aid::gfp]) X; ift-122(tm2878) IV; mks-5(tm3100) II* | IFT-54::GFP + *mks-5* + *ift-122* | *this study* |

**Table S2.** List of primers used in this study.

| | Screen | Primer ID | Sequence (5'-3') |
|---|---|---|---|
| Dynein-2 complex | DHC-2 | oTD54 (FW) | ACAGGTGGAGTGTATTTAATGAG |
| | | oTD2 (RV) | ACAACACGAAAGACGTTGGCTG |
| | WDR-60 | oTD41 (RV) | CAAATGCCACAAGATACGGAC |
| | | oTD42 (FW) | TCATAACATACTGGGATTTGGG |
| IFT-B Complex | IFT-54 | oTD615 (FW) | AGATAGAGGAGCTTTGGTG |
| | | oTD616 (RV) | TTAATCTGGGTTTTCCTGTG |
| | IFT-74 | oTD5 (FW) | CACGAGTATGACTCACAAGGAG |
| | | oTD6 (RV) | CGGAAAGGGTGCTTCATACTTG |
| IFT-A Complex | *ift-122 (tm2878)* | oTD606 (FW) | GCAATTTGAACTGGATGGTC |
| | | oTD571 (RV) | TACACTCTGGTTAGGGTCAC |
| | *ift-140 (tm3433)* | oTD161 (FW) | TCTTCGTAGTATCTTGCAGCG |
| | | oTD162 (RV) | TACAACTTCAAGGAATGCAGC |
| | *ift-144 (gk678)* | oTD413 (FW) | CGGTTACCAAGGCAACAAC |
| | | oTD414 (RV) | CGCATTGGTCATGAAGCTC |
| Transition Zone components | *nphp-4 (tm925)* | oTD153 (FW) | GCCATTCTTGGATCCGCATA |
| | | oTD154 (RV) | CCGGCCAGTTGAAATGAAAC |
| | *mks-5 (tm3100)* | oTD393 (FW) | TTCAATAGCAGATTTCGCCG |
| | | oTD394 (RV) | AATCACATTCCTCTTGCAGC |
| Ciliary Base | SPD-5 | oTD752 (FW) | CTTGTTTCAGAAAACTTCGCG |
| | | oTD753 (RV) | CCTTAACTCTCTTCTGCGACG |
| Ciliary Evs | TSP-6 | oTD708 (FW) | ATTTACGTCGCCTTCGTTGG |
| | | oTD709 (RV) | GCAGTTTACTCGTAGGGCGA |
| Ciliary Axoneme | MosSCI Chr IV cxTi10882 | oTD271 (RV) | TTTACAAGGACTTGGATAAATTGG |
| | | oTD320 (FW) | CAGGAGAGCAAGGACCAAAG |
| | | oTD319 (RV) | GCCAAATGCCATAGTCAATGG |
